# Leveraging DNA methylation quantitative trait loci to characterize the relationship between methylomic variation, gene expression and complex traits

**DOI:** 10.1101/297176

**Authors:** Eilis Hannon, Tyler J Gorrie-Stone, Melissa C Smart, Joe Burrage, Amanda Hughes, Yanchun Bao, Meena Kumari, Leonard C Schalkwyk, Jonathan Mill

## Abstract

Characterizing the complex relationship between genetic, epigenetic and transcriptomic variation has the potential to increase understanding about the mechanisms underpinning health and disease phenotypes. In this study, we describe the most comprehensive analysis of common genetic variation on DNA methylation (DNAm) to date, using the Illumina EPIC array to profile samples from the UK Household Longitudinal study. We identified 12,689,548 significant DNA methylation quantitative trait loci (mQTL) associations (P < 6.52x10^-14^) occurring between 2,907,234 genetic variants and 93,268 DNAm sites, including a large number not identified using previous DNAm-profiling methods. We demonstrate the utility of these data for interpreting the functional consequences of common genetic variation associated with > 60 human traits, using <u>S</u>ummary data–based <u>M</u>endelian <u>R</u>andomization (SMR) to identify 1,662 pleiotropic associations between 36 complex traits and 1,246 DNAm sites. We also use SMR to characterize the relationship between DNAm and gene expression, identifying 6,798 pleiotropic associations between 5,420 DNAm sites and the transcription of 1,702 genes. Our mQTL database and SMR results are available via a searchable online database (http://www.epigenomicslab.com/online-data-resources/) as a resource to the research community.

## INTRODUCTION

DNA methylation (DNAm) is an epigenetic modification to cytosine that is involved in mediating the developmental regulation of gene expression and function, and transcriptional processes including genomic imprinting and X-chromosome inactivation (Bird 2002; Jones 2012). Although often regarded as a mechanism of transcriptional repression, the relationship between DNAm and gene expression is highly complex and not fully understood (Wagner et al. 2014). Gene body DNAm, for example, is often associated with active expression (Yang et al. 2014), and also influences other transcriptional processes including alternative splicing and promoter usage (Maunakea et al. 2010). There is growing interest in the role of DNAm in disease, with recent epigenome-wide association studies (EWAS) identifying robust associations between variable DNAm and cancer (Baylin and Jones 2016) and a diverse range of other complex phenotypes including rheumatoid arthritis (Liu et al. 2013), body mass index (Wahl et al. 2017), schizophrenia (Hannon et al. 2016) and Alzheimer’s disease (Lunnon et al. 2014). Characterizing the complex relationship between genetic, epigenetic and transcriptomic variation will increase understanding about the mechanisms underpinning health and disease phenotypes. Twin and family studies have demonstrated that population-level variation in DNAm is under considerable genetic control, although these effects vary across genomic loci, developmental stages, and different cell- and tissue-types (Bell et al. 2012; Grundberg et al. 2013; McRae et al. 2014; Hannon et al. 2015; van Dongen et al. 2016). Studies in a variety of tissues, including brain, whole blood, pancreatic islet cells, and adipose tissue, have identified widespread associations between common DNA sequence variants and DNAm (Gibbs et al. 2010; Drong et al. 2013; Gamazon et al. 2013; Olsson et al. 2014; Hannon et al. 2015; Gaunt et al. 2016). These DNAm quantitative trait loci (mQTLs) are primarily *cis-*acting, are enriched in regulatory chromatin domains and transcription factor binding sites, and have been shown to colocalize with gene expression quantitative trait loci (eQTLs)(Gutierrez-Arcelus et al. 2013; Wagner et al. 2014; Hannon et al. 2015).

There is considerable interest in using mQTLs, along with other types of molecular QTL, to interpret the functional consequences of common genetic variation associated with human traits, especially as the actual gene(s) involved in mediating phenotypic variation are not necessarily the most proximal to the lead SNPs identified in genome-wide association studies (GWAS). Of note, GWAS variants are enriched in enhancers and regions of open chromatin (Ernst et al. 2011; Schaub et al. 2012), reinforcing the hypothesis that most common genetic risk factors influence gene regulation rather than directly affecting the coding sequences of transcribed proteins (Maurano et al. 2012). Importantly, evidence for the co-localization of genetic variants associated with both phenotypic and regulatory variation is not sufficient to show that the overlapping association signals are causally related; additional analytical steps are needed to distinguish pleiotropic effects - i.e. where the same variant is influencing both outcomes, although not necessarily dependently - from those that are an artefact of linkage disequilibrium (LD). We recently extended the use of one approach - <u>S</u>ummary data–based <u>M</u>endelian <u>R</u>andomization (SMR), which was initially used in conjunction with eQTL data (Zhu et al. 2016) - to prioritize genes for GWAS-nominated loci using mQTL data (Hannon et al. 2017).

Building on our previous work, we have performed the most comprehensive analysis of genetic effects on DNAm yet undertaken. We used the Illumina EPIC array and imputed SNP data to identify mQTLs associated with variable DNAm at ~850,000 sites across the genome in samples from the Understanding Society UK Household Longitudinal study (UKHLS) (n = 1,111). These mQTL were used within the SMR framework to refine genetic association data from publically-available GWAS datasets in order to prioritize genes involved in 63 complex traits and diseases. We subsequently used the SMR approach to identify pleiotropic relationships between DNAm and variable gene expression using publically-available whole blood gene eQTL data. Our mQTL database and SMR results are available via a searchable online database (http://www.epigenomicslab.com/online-data-resources/) as a resource to the research community.

## RESULTS

### Novel mQTL associations identified using the Illumina EPIC array

An overview of our study design is presented in **Supplementary Figure 1**. We tested 5,210,475 imputed genetic variants against the 766,714 DNAm sites on the Illumina EPIC array passing our stringent QC criteria (see **Methods**). We identified 12,689,548 significant mQTL associations at a conservative Bonferroni-corrected threshold (P < 6.52x10^-14^) occurring between 2,907,234 genetic variants and 93,268 DNAm sites (**Supplementary Table 1; Figure 1A**) with a mean percentage point change in DNAm per allele across all mQTL-associated sites of 3.46% (SD = 3.01%). Existing mQTL databases have been almost exclusively generated using the Illumina 450K array; more than half of the DNAm sites (n = 48,099, 51.6%; **Supplementary Table 2**) we identify as being associated with genetic variation using the Illumina EPIC array involve novel content not previously interrogated (**Supplementary Figure 2**). Importantly, these novel mQTL associations are annotated to 5,172 genes not included in mQTL databases generated using the Illumina 450K array (**Supplementary Figure 3)**. DNAm sites associated with genetic variation are associated with a median of 65 mQTLs (SD = 238), probably reflecting linkage disequilibrium (LD) relationships between proximal variants. In contrast, each mQTL variant is associated with a median of two (SD = 6.37) DNAm sites, with the majority of mQTL SNPs (n = 1,003,238, 34.5%) being associated with DNAm at only a single site (**Supplementary Figure 4**). We performed LD clumping of the results for each DNAm site to identify the number of *independent* associations for each DNAm site (see **Methods**); this process reduced the number of mQTL associations (P < 6.52x10^-14^) to 161,761 (1.27% of the total number of unclumped significant mQTL associations), with a median of 1 (SD = 1.61) mQTL variant associated with each DNAm site (**Supplementary Figure 5**). At a more relaxed significance threshold (P < 1x10^-10^) we identified a total of 17,051,673 mQTL associations between 3,281,391 genetic variants and 114,595 DNAm sites; these results are available in a searchable database at http://www.epigenomicslab.com/online-data-resources/.

**Figure 1.**
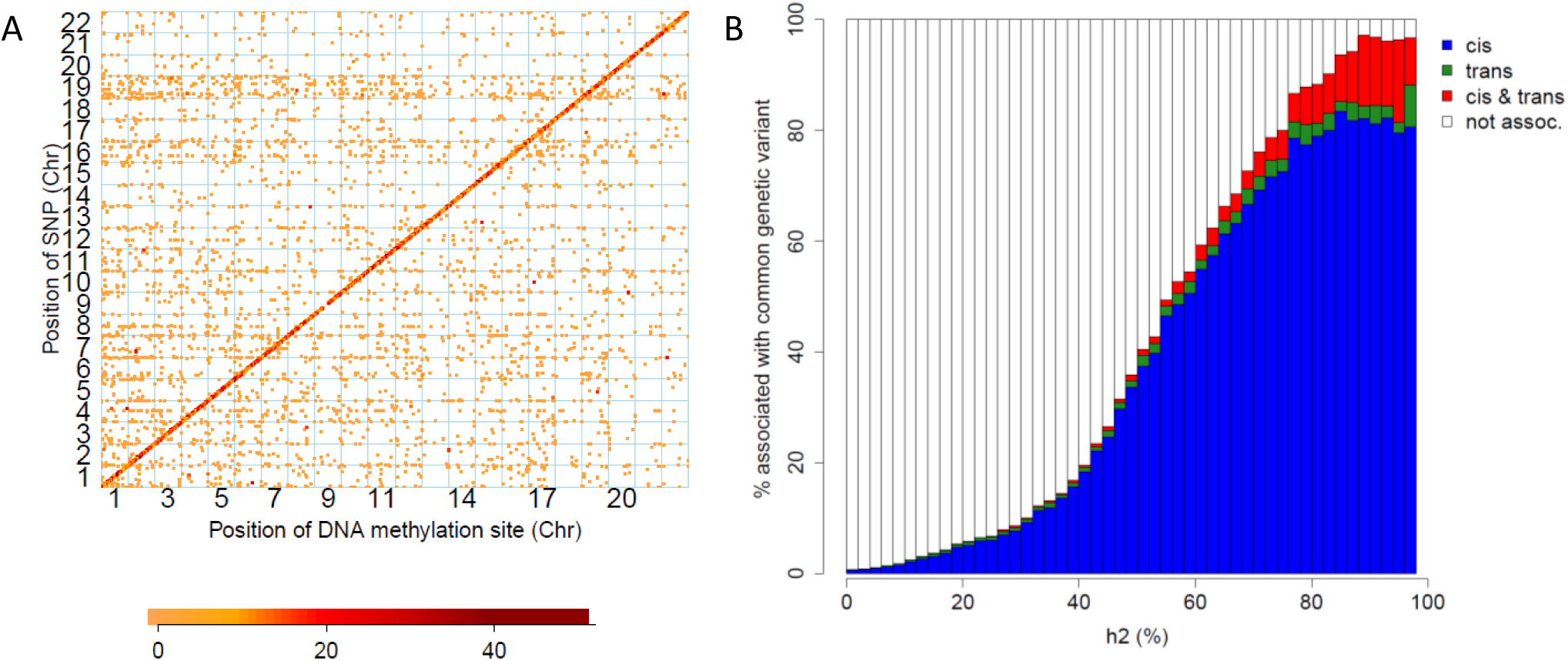
DNA methylation quantitative trait loci are predominantly *cis*-acting and enriched in sites known to be highly heritable. **A)** The genomic distribution of Bonferroni significant (*P* = 6.52x10^-14^) mQTLs in whole blood, where the position on the x-axis indicates the location of Illumina EPIC array probes and the position on the y-axis indicates the location of genetic variants. The color of the point corresponds to the difference in DNA methylation per allele compared to the reference allele, with the largest effects plotted in dark red. A clear positive diagonal can be observed demonstrating that the majority of mQTLs are associated with genotype in *cis*. **B)** A barplot of the percentage of DNA methylation sites associated with common genetic variation grouped by previous reported estimates of heritability (% variation explained by additive genetic factors for each site taken from (van Dongen et al. 2016)van Dongen et al.). Each barplot demonstrates the percentage of DNA methylation sites with Bonferroni significant genetic effects in *cis* only (blue), *trans* only (green), both *cis* and *trans* (red) and no significant genetic effects (white).

**Figure 2.**
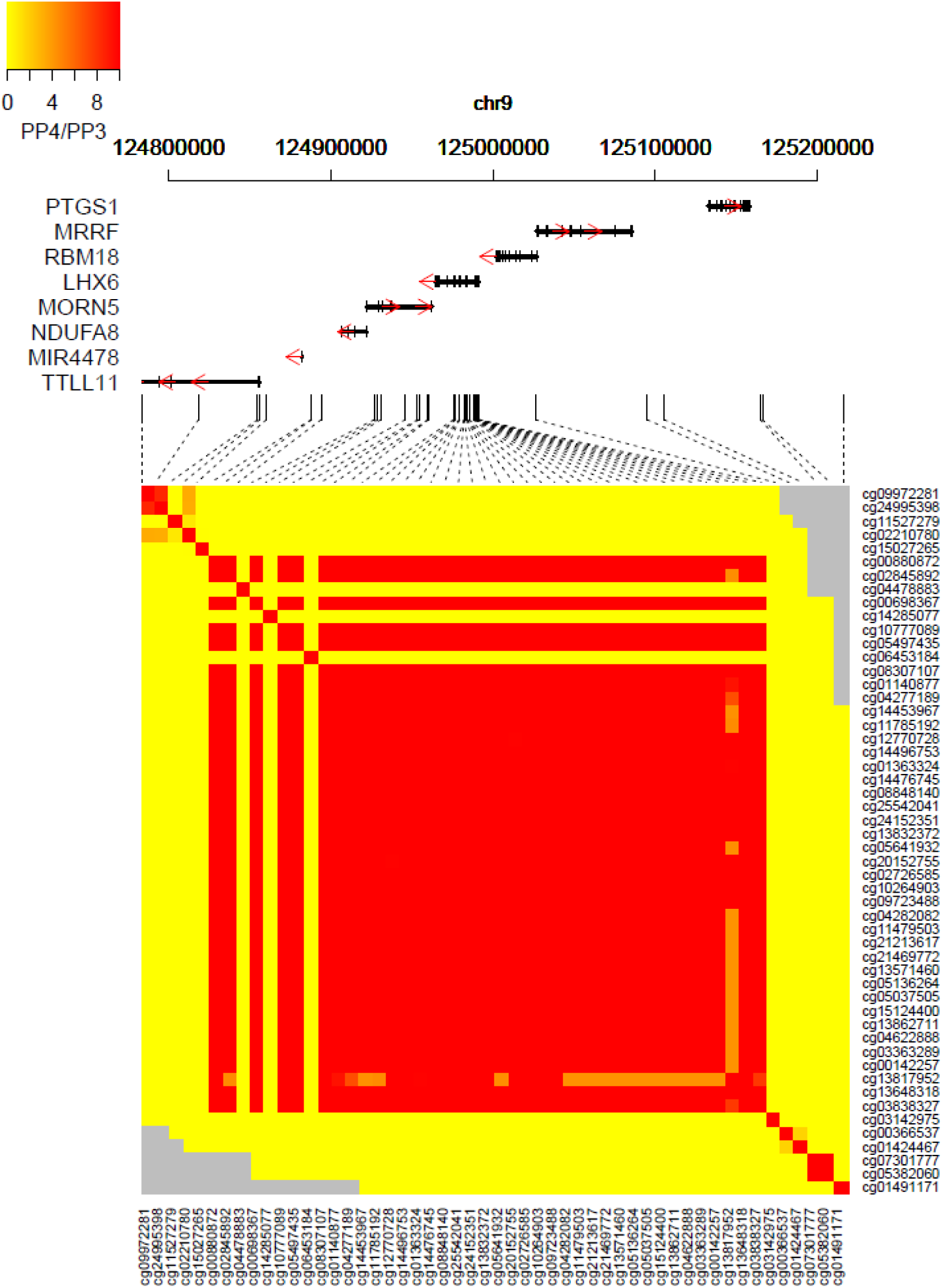
Shared genetic architecture between neighbouring DNA methylation sites. Heatmap of Bayesian co-localisation results for all pairs of DNA methylation sites with at least one significant mQTL (P < 1x10^-10^) in a genomic region on chromosome 9 (chr9:124783559-125216341). Columns and rows represent individual DNA methylation sites (ordered by genomic location). The color of each square indicates the strength of the evidence for a shared genetic signal (from yellow (weak) to red (strong)) calculated as the ratio of the posterior probabilities that they share the same causal variant (PP4) compared to two distinct causal variants (PP3). The ratio was bounded to a maximum value of 10; gray indicates pairs of DNA methylation sites that were not tested for co-localization.

**Figure 3.**
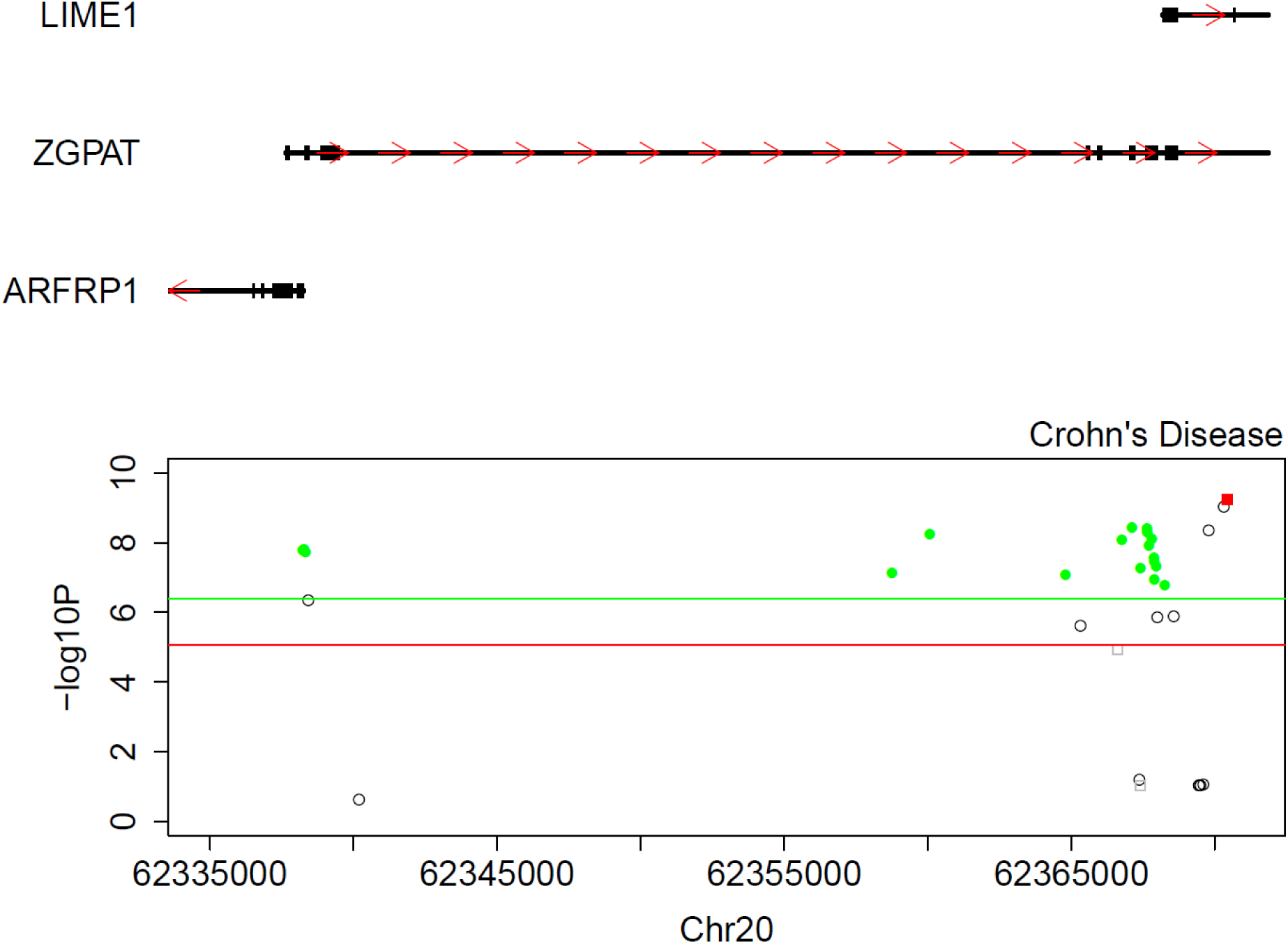
<u>S</u>ummary data–based <u>M</u>endelian <u>R</u>andomization (SMR) analysis using quantitative trait loci associated with DNA methylation (mQTL) and gene expression (eQTL) implicates a role for *LIME1* in Crohn’s disease. Shown is a genomic region on chromosome 20 (chr20:62335000-62371000) identified in a recent GWAS of Crohn’s disease performed by (Liu et al. 2015). Genes located in this region are shown at the top, with exons indicated by thicker bars and the red arrows indicating the direction of transcription. The scatterplot depicts the –log10 *P* value (y-axis) against genomic location (x-axis) from the SMR analysis (where circles represent Illumina EPIC array DNA methylation sites and squares represent gene expression probes, with solid green and red highlighting those with non-significant HEIDI test for DNA methylation and gene expression respectively). The green and red horizontal lines represent the multiple testing corrected threshold for the SMR test using mQTL and eQTL respectively.

**Figure 4.**
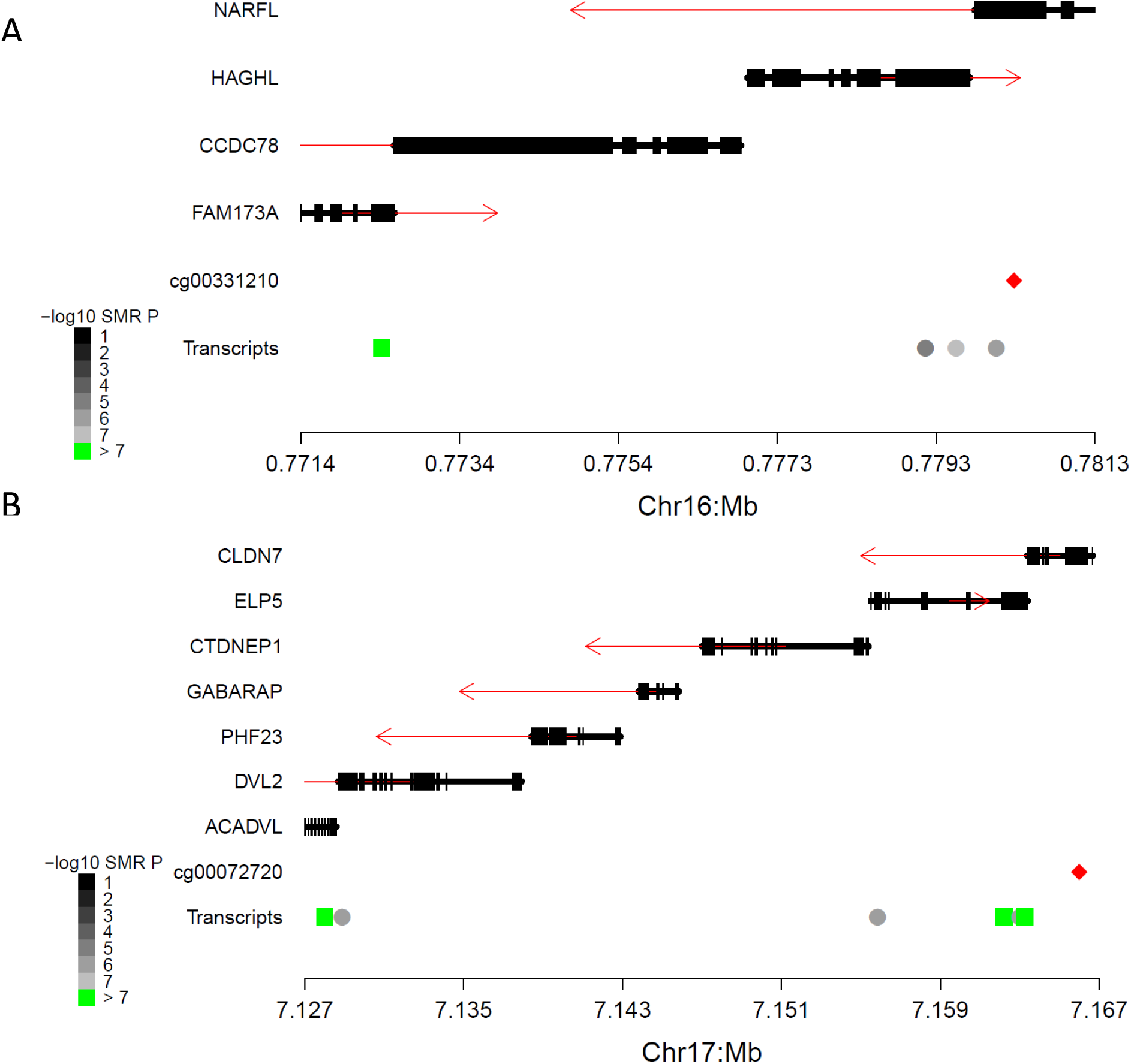
Regional plots demonstrating the complex relationship between gene expression and DNA methylation identified using SMR. Shown is an example of **A)** a DNA methylation site (cg00331210) that is associated with expression of a gene (FAM173A) that is not the most proximal to it and **B)** a DNA methylation site (cg00072720) associated with the expression of multiple genes (CLDN7 and ELP5). Each plot contains a gene track where red arrows indicate the direction of transcription and a red diamond indicates the position of the pleiotropically significant DNA methylation site. Circles and squares indicate the location of the gene expression probes that that DNA methylation sites were tested against where the colour indicates the significance level of the SMR test (black to gray) with green indicating significant associations (P < 1.02x10^-7^). For significant associations squares indicate tests with non-significant heterogeneity (P > 0.05) that are indicative of pleiotropic associations.

### mQTL associations predominantly occur in cis and influence DNAm at sites known to be influenced by heritable factors

Consistent with the results of previous studies, we found that the majority of mQTL associations (n = 11,679,376; 92%) occur in *cis*, defined as situations where the distance between mQTL SNP and DNAm site is ≤ 500kb (Gibbs et al. 2010; Drong et al. 2013; Olsson et al. 2014; Hannon et al. 2015) (**Figure 1A**). *Cis* mQTL variants are typically associated with larger effects on DNAm than those acting in *trans* (average *cis* effect = 3.48% change in DNAm per allele, average *trans* effect = 3.26% change in DNAm per allele; Mann Whitney P < 2.23x10^-308^) (**Supplementary Figure 6)**. Furthermore, amongst *cis* mQTL associations, both significance and effect size increase as the distance between the genetic variant and DNAm site decreases (**Supplementary Figure 7**). Compared to the distribution of all tested DNAm sites, those associated with at least one mQTL variant (correcting for the number of tests performed (see **Methods**), P < 6.52x10^-14^) are significantly enriched in intergenic regions and less likely to be located within both gene bodies (Chi square test: P < 2.23x10^-308^; **Supplementary Figure 8; Supplementary Table 3A**) and CpG islands (Chi square test P < 2.23x10^-308^; **Supplementary Figure 9; Supplementary Table 3B**). We used quantitative genetic data from a study of DNAm in monozygotic and dizygotic twins (van Dongen et al. 2016) to show that DNAm at sites associated with at least one mQTL variant is more strongly influenced by heritable (additive genetic) factors compared to all DNAm sites tested (mQTL sites: median heritability (h^2^) = 55% (SD = 22.8%); all DNAm sites: median h^2^ = 12% (SD = 20.5%); Mann Whitney P < 2.23x10^-308^; **Supplementary Figure 10**). Overall, the proportion of sites at which DNAm is associated with an mQTL variant increases as a function of the estimated additive genetic influence derived from twin analyses (**Figure 1B**). Interestingly, there is no significant difference in the contribution of additive genetic effects to variance in DNAm at sites associated with *cis* (median h^2^ = 56%; SD = 22.5%) and *trans* (median h^2^ = 57%; SD = 26.9%) mQTL variants (Mann-Whitney P = 0.910).

### Proximal DNA methylation sites share genetic associations

Similar to the LD relationships that exist between proximal genetic variants, DNAm levels are often correlated between proximally located DNAm sites (Bell et al. 2012; Liu et al. 2014). To further characterize the genetic architecture of DNA methylation, we investigated whether shared genetic effects on multiple DNAm sites underlies this regional correlation structure. Although genetic variants are often associated with variation at multiple DNAm sites (**Supplementary Figure 4**), this does not establish a shared genetic effect; shared genetic signals influencing a pair of DNAm sites may result from two distinct causal genetic variants that are in strong LD. To formally test whether neighbouring DNAm sites are influenced by the same causal variant, we used a Bayesian co-localization approach (Giambartolomei et al. 2014) to interrogate all pairs of DNAm sites characterized as being located within 250kb of each other and associated with at least one significant mQTL variant (P < 1x10^-10^). Our analyses assessed 3,535,812 pairs of DNAm sites with a median distance between DNAm sites of 110,493bp (SD = 74,448 bp), comparing the pattern of mQTL associations for both DNAm sites to test whether they index an association with either the same causal variant or two distinct causal variants. We found that the posterior probabilities for virtually all (n = 3,520,781 (99.6%), median distance of 110,319bp (SD = 74,440bp)) supported a co-localized association within the same genomic region (PP_3_+PP_4_ > 0.99). Of these, 281,898 pairs (8%) had sufficient support for both DNAm sites being associated with the same causal mQTL variant (PP_3_ + PP_4_ > 0.99 & PP_4_/PP_3_ > 1; **Supplementary Table 4**), with 234,460 pairs (6.6%) having ‘convincing’ evidence (PP_3_ + PP_4_ > 0.99 & PP_4_/PP_3_ > 5) for co-localization of the same mQTL association according to the criteria of Guo and colleagues (Guo et al. 2015). DNAm sites that shared genetic effects with at least one other DNAm site co-localize with a median of three other DNAm sites, indicating a complex relationship between genetic variation and DNAm in *cis*. **Figure 2**, for example, demonstrates a broad genomic region (> 400 kilobases) on chromosome 9 where 38 DNAm sites - spanning seven genes - have a common underlying genetic signal. Of note, these DNAm sites are not contiguous; a small number of genetically-mediated DNAm sites located within this region do not share the same mQTL signal. Pairs of DNAm sites with a shared causal mQTL variant are enriched for concordant directions of effect (71.2% pairs with positive correlations vs 28.8% pairs with negative correlations, binomial test P = 1.48x10^-323^; **Supplementary Figure 11**). Furthermore, these pairs are located relatively close together (median distance between convincing co-localized pairs = 12,394 bp (SD = 53,451bp)), with evidence that the shared genetic architecture is structured around annotated genomic features. Co-localized pairs of DNAm sites are significantly more likely to be annotated to the same gene (OR = 6.08, Fisher’s test P < 2.23x10^-308^) or CpG island (OR = 1.54, Fisher’s test P < 2.23x10^-308^) compared to non-co-localized pairs. Where pairs of DNAm sites with a shared genetic signal are annotated to the same gene, they are nominally less likely to be annotated to the same feature compared to pairs of DNAm sites annotated to different genes (OR = 0.956, Fisher’s test P = 2.52x10^-7^) suggesting that where genetic variation influences DNAm at multiple sites across a gene they do not necessarily cluster by genic feature and can be located anywhere from the transcription start site to the end of the last exon. DNAm is more likely to be positively correlated between pairs of co-localized sites annotated to the same gene compared to pairs of sites annotated to different genes (OR = 1.85, Fisher’s P < 2.23x10^-308^), a result driven predominantly by pairs of DNAm sites annotated to the same feature within that gene (OR = 1.57, Fisher’s test P = 3.41 x10^-135^) rather than those annotated to different features within a gene. Finally, pairs of DNAm sites with shared genetic effects annotated to the same genic feature, although not necessarily the same gene, are more likely to be positively correlated compared to pairs annotated to different genic features (OR = 1.73, Fisher’s P < 2.23x10^-308^; **Supplementary Figure 12**).

### DNAm quantitative trait loci have utility for refining GWAS signals for complex traits

Genetic variants identified in GWAS analyses rarely index protein-coding changes. Instead they are hypothesized to influence gene regulation, being enriched in regulatory motifs including enhancers and regions of open chromatin (Maurano et al. 2012; Schaub et al. 2012). There is considerable interest in using regulatory QTLs to refine genetic association signals and prioritize potentially causal genes within the extended genomic regions identified in GWAS (Hannon et al. 2015; Zhu et al. 2016; Hauberg et al. 2017; Mancuso et al. 2017). We next extended our previous application of the SMR approach (Hannon et al. 2017) to test 126,457 DNAm sites against 63 complex phenotypes with GWAS data (**Supplementary Table 5**). The first stage of the SMR approach uses the most significantly associated mQTL SNP associated with DNAm at a specific site - that has also been tested in the GWAS dataset - as an instrumental variable, implementing a two-step least squares (2SLS) approach to compare the estimated associations. Using this approach, we identified 5,848 associations (*P* < 3.95x10^-7^ corrected for 126,457 DNAm sites) between 40 complex traits and 5,849 unique DNAm sites (**Supplemental File 1**). Because the associations identified in this way potentially reflect two highly-correlated but different causal variants for the GWAS trait and DNAm, the second stage of the SMR method repeats the analysis with alternative mQTL SNPs as the instrument. If there is a single causal variant associated with both phenotype and DNAm, the association statistics will be identical regardless of the selected instrument. In contrast, if there are two separate causal variants, each correlated with the instrument, there will be variation in the results. To distinguish between these scenarios, we applied the heterogeneity in dependent instruments (HEIDI) test to select associations with non-significant heterogeneity (HEIDI *P* > 0.05), identifying a refined set of 1,662 associations between 36 complex traits and 1,246 DNAm sites (**Supplementary Table 6**).

As the power of the SMR approach to detect pleiotropic associations reflects, in part, the power of the initial complex trait GWAS, it is unsurprising that the highest number of SMR associations were found for traits characterized by the largest number of GWAS signals such as height (423 significant GWAS loci, 506 SMR pleiotropic associations)(Wood et al. 2014) and inflammatory bowel disease (168 significant GWAS variants, 127 SMR pleiotropic associations)(Liu et al. 2015). In contrast, no SMR associations were found for traits with few or no genome-wide significant SNPs including parental age at death (0-1 significant GWAS variants)(Pilling et al. 2016), insulin secretion rate (no significant GWAS variants)(Wood et al. 2017) and whether a person has ever smoked (no significant GWAS variants)(Tobacco and Genetics Consortium 2010). We compared our SMR results to those obtained using our previous mQTL dataset - generated using a smaller number of samples - observing high rates of replication for loci that were tested in both analyses. As our previous SMR analysis was based on a subset of 43 traits and the reduced content of the Illumina 450K array, 842 pleiotropic associations reported in the current analysis were taken forward for replication; DNAm at 519 (33.0%) of these was associated with an mQTL variant and therefore were tested in our previous SMR study, with 268 (51.6%) characterized by significant pleiotropic association in both studies. Furthermore, the vast majority of associations tested in both datasets (516; 99.4%) were in the same direction, significantly more than expected by chance (sign test P = 2.72x10^-149^ **Supplementary Figure 13**), suggesting that there are many additional true signals in those that did not meet the stringent criteria for significance used in both studies.

In order to prioritise genes for each complex trait we characterized the genic location of associated DNAm sites. 1,269 (76.3%) of the identified pleiotropic associations involve DNAm sites located within a gene or less than 1500bp from the transcription start site, representing a significantly higher rate compared to all DNAm sites tested in our SMR analysis (OR = 1.64, Fisher’s test P = 1.12x10^-18^). To further explore these 786 pleiotropic associations – occurring between 577 genes and 32 complex traits - we extended our SMR analyses to incorporate a publically-available whole blood gene expression quantitative trait loci (eQTL; n = 5,311 individuals) dataset (Westra et al. 2013). Expression of 232 (40.2%) of our identified genes was significantly associated with an eQTL variant and these were used to test for pleiotropic associations between gene expression level and the GWAS trait. These analyses provided additional support for 138 of the pleiotropic associations identified using mQTL data, supporting a relationship between 33 genes and 17 complex traits (**Supplementary Table 7**). **Figure 3**, for example, highlights an association between *LIME1* and Crohn’s Disease characterized by significantly pleiotropic relationships with multiple DNAm sites around the transcription start site and also expression of *LIME1*.

### Pleiotropic associations between DNAm and gene expression

Although it is widely hypothesized that DNAm influences gene expression, its relationship with transcriptional activity is not fully understood. DNAm across CpG-rich promoter regions, for example, is often assumed to repress gene expression via the blockage of transcription-factor binding and the attraction of methyl-binding proteins (Bogdanović and Veenstra 2009). DNAm in the gene body, in contrast, is hypothesized to be a marker of active gene transcription (Ball et al. 2009; Maunakea et al. 2010), potentially playing a role in regulating alternative splicing and isoform diversity. To identify associations between DNAm and gene expression we applied the SMR approach to DNAm sites located within a megabase of a gene expression probe included in the eQTL dataset generated by Westra and colleagues (Westra et al. 2013). In total, we tested 488,342 pairs, exploring relationships between 96,694 DNAm sites and 4,721 gene expression probes annotated to 4,049 genes (**Supplementary Figure 14**). On average, each DNAm site was tested against a median of four expression probes (SD = 4.06) mapping to a median of three genes (SD = 3.45). In contrast, each expression probe was tested against a median of 85 DNAm sites (SD = 72.8). Of these, 40,404 pairs (8.27%) - comprising 22,007 (22.8%) DNAm sites and 4,201 (89.0%) expression probes mapping to 3,628 (89.6%) genes - were characterized by a significant SMR result (significance threshold corrected for the number of DNAm sites and gene expression probe pairs tested = P < 1.02x10^-7^). 6,798 of these significant SMR pairs - comprising 5,420 (5.61%) DNAm sites and 1,913 (40.5%) expression probes mapping to 1,702 (42.0%) genes - also had a HEIDI P > 0.05 (**Supplementary Table 8; Supplementary Figure 14**). These results suggest that while expression of a large proportion of genes is associated with DNAm sites, not all DNAm sites are associated with gene expression in *cis*.

The majority of significant gene expression probes (n = 1,192; 62.3%) are associated with a median of two DNAm sites (SD = 4.17 sites) spanning a median distance of 66,846bp (SD = 163,802bp) at a median density of 19,959bp (SD = 67,498bp) between sites. Interestingly, DNAm sites pleiotropically associated with gene expression are enriched in the gene body and transcription start sites of genes and depleted intergenically (Chi square test P = 7.08x10^-133^; **Supplementary Figure 15; Supplementary Table 9**). We identified a small but significant enrichment of scenarios where DNAm is negatively associated with gene expression at sites located in the 5’UTR (P = 0.00108), TSS200 (P = 6.38x10^-7^), TSS1500 (P = 5.82x10^-11^) and 1^st^ exon (P = 6.19x10^-5^), consistent with the hypothesis that promoter DNAm often represses gene expression (**Supplementary Figure 16**).

### Using QTL data to refine the genic annotation associated with DNAm sites

A key challenge in epigenetic epidemiology relates to how DNAm sites are assigned to genes in order to facilitate the biological interpretation of significant EWAS associations. DNAm sites are usually annotated to specific genes based on proximity, although the extent to which this approach is valid for inferring downstream transcriptional effects is not known. Among the identified pleiotropic associations between DNAm and gene expression we selected instances where the DNAm site is not intergenic - i.e. < 1500bp from the transcription start site of a gene (n = 5,593 (82.3%)) - finding that these were annotated to the same gene whose expression level they were associated with at a much higher rate compared to DNAm sites significantly associated with expression levels at another gene (OR = 9.67; Fisher’s test P < 2.23x10^-308^). Of the 5,460 DNAm sites significantly associated with expression of at least one gene, 1,790 (32.8%) were associated with the gene they were annotated to, although 276 (5.05%) of these were also associated to an additional gene and 2,686 (50.0%) were associated with a different gene. Of note, not all CpGs were tested against the gene they were annotated to because the gene lacked a significant eQTL; this was the case for the majority of DNAm sites (n = 2,701; 80.4%) identified as associated with a gene other than the one they were annotated to. Of particular interest are the 944 (18.3%) sites classed as intergenic which are associated with gene expression, potentially providing novel gene annotations for interpreting the results of EWAS analyses. Overall, although the proximity-based annotation of DNAm sites appears to be appropriate in many instances, we identified notable exceptions. For example, **Figure 4A** shows that the DNAm site cg00331210, located within the body of *NARFL* on chromosome 16, is not associated with expression of that gene, but with the *FAM173A* gene, which is located 7.9 kilobases away. Likewise, **Figure 4B** shows that the DNAm site cg00072720, located within the gene body of *CLDN7*, is not associated with expression of that gene but two other genes (*ACADVL* and *ELP5/C17ORF81*) on chromosome 17.

## DISCUSSION

In this study we present the most comprehensive assessment of the genetic architecture of DNAm to date, identifying associations between common genetic variants and specific DNAm sites (mQTLs), using the Illumina EPIC array. We utilized our novel database of mQTL associations to characterize genetic influences on individual and proximally-located DNAm sites. We show that there are many instances of shared genetic signals on neighbouring DNAm sites and that these associations are structured around both genes and CpG islands. Moreover, we report that these shared genetic effects on DNAm are generally associated with positive correlations between the DNAm sites. This has implications for studies of trait-associated differentially methylated regions (DMRs), as it suggests that associations with phenotypic variation could be genetically mediated.

We extended our previous work prioritizing genes in GWAS-nominated regions (Hannon et al. 2017), finding robust agreement with our previous SMR findings (using mQTLs identified with the Illumina 450K array) for shared content using independent datasets. The additional content present on the EPIC array, however, enabled us to identify novel gene-trait associations not detected using the older array technology, increasing the potential yield of biological information. Finally, we use these data to explore the relationship between DNAm and gene expression using genetic instruments rather than correlations to infer associations between specific DNAm sites and genes. Although most DNAm sites associated with gene expression were found to be located within the gene body or close to the transcription start site, there are many relationships that challenge the commonly used genic annotation based solely on physical proximity. Furthermore, while the expression of most genes is associated with one or more DNAm sites, not all DNAm sites are associated with gene expression, implying that variable DNAm does not always have an effect on gene expression. Although we could only test for associations between DNAm sites with significant mQTLs and the expression of genes with a significant eQTL, our results provide a potentially effective method for annotating results from EWAS, particularly where the influence of DNAm on gene expression is hypothesized and candidates are taken forward for transcriptional analysis.

Our study has a number of important limitations. The analyses presented here are based on an unrelated subset participants from the UKHLS; although these represent a large sample (> 1,000) of European ancestry with a broad age range, the extent to which our results are applicable to other ethnic groups characterized by a different genetic architecture is not known. Despite using the most comprehensive, high-throughput technology for profiling DNAm across the genome (the Illumina EPIC array), our study only assayed a small proportion of the total number DNAm sites with sparse coverage of regulatory features that are often represented by a single DNAm site (Pidsley et al. 2016). Moreover, DNAm was profiled in whole blood, which potentially limits the interpretation of candidate disease genes where the presumed tissue of interest is not blood. Given the tissue-specific nature of some mQTL and eQTL effects these associations should be confirmed in additional disease relevant tissues and cell types. Finally, although Mendelian Randomisation is proposed as a methodology for quantifying causal relationships between variables, it is reliant on a number of key assumptions (Smith and Ebrahim 2003) all of which also apply to SMR. We are therefore careful in our use of terminology and refrain from describing our associations as “causal”, especially because the SMR approach is unable to distinguish two causal variants in approximately perfect LD from one causal variant (Zhu et al. 2016); instead we refer to these as “pleiotropic” associations. Furthermore, it is possible that variants may act through mechanisms such as horizontal pleiotropy.

Taken together, our results add to an increasing body of evidence showing that genetic influences on DNA methylation are widespread across the genome. We show that these relationships may be integrated with the results from GWAS of complex traits and genetic studies of gene expression in order to improve our understanding about the interplay between gene regulation and expression, facilitating the prioritization of candidate genes implicated in disease aetiology.

## METHODS

### Sample description

The British Household Panel Survey (BHPS) began in 1991, and in 2010 was incorporated into the larger UK Household Longitudinal Study(Knies 2015) (UKHLS; also known as Understanding Society (https://www.understandingsociety.ac.uk)) which is a longitudinal panel survey of 40,000 UK households from England, Scotland, Wales and Northern Ireland. Since 1991 annual interviews have collected sociodemographic information, and in 2011-12, biomedical measures and blood samples for BHPS participants were collected at a nurse visit in the participant’s home. Respondents were eligible to give a blood sample if they had taken part in the previous main interview in English, were aged 16+, lived in England, Wales or Scotland, were not pregnant, and met other conditions detailed in the user guide (Benzeval M. et al. 2014). For each participant, non-fasting blood samples were collected through venepuncture; these were subsequently centrifuged to separate plasma and serum, aliquoted and frozen at −80 °C. DNA has been extracted and stored for genetic and epigenetic analyses.

### Genome-wide quantification of DNAm

DNAm was profiled in DNA extracted from whole blood for 1,193 individuals aged from 28 to 98 who were eligible for and consented to both blood sampling and genetic analysis, had been present at all annual interviews between 1999 and 2011, and whose time between blood sample collection and processing did not exceed 3 days. Eligibility requirements for genetic analyses meant that the epigenetic sample was restricted to participants of white ethnicity. 500ng of DNA from each sample was treated with sodium bisulfite, using the EZ-96 DNA methylation-Gold kit (Zymo Research, CA, USA). DNAm was quantified using the Illumina Infinium HumanMethylationEPIC BeadChip (Illumina Inc, CA, USA) run on an Illumina iScan System (Illumina, CA, USA) using the manufacturers’ standard protocol. Samples were randomly assigned to chips and plates to minimise batch effects. In addition, a fully methylated control (CpG Methylated HeLa Genomic DNA; New England BioLabs, MA, USA) was included in a random position on each plate to facilitate sample tracking, resolve experimental inconsistencies and confirm data quality.

### DNAm data preprocessing

Raw signal intensities were imported from idats into the R statistical environment(R Development Core Team 2008) and converted into beta values using the bigmelon package (Gorrie Stone et al. submitted). Data was processed through a standard pipeline and included the following steps: outlier detection, confirmation of complete bisulphite conversion, estimating age from the data (Horvath 2013) and comparing with reported age visualisation of principal components. Data were normalized using the dasen function from the wateRmelon package(Pidsley et al. 2013). Samples that were dramatically altered as a result of normalisation were excluded by assessing the difference between normalized and raw data and removing those with a root mean square and standard deviation > 0.05. were then filtered to exclude samples and then DNA methylation sites with > 1% of sites or samples with detection p value > 0.05, finally DNA methylation sites with a bead count < 3 were also excluded. The data were then re-normalised with the dasen function. The final dataset included 857, 071 DNA methylation sites and 1, 175 individuals for analysis.

### Annotation of DNAm sites

The genomic location of DNAm sites along with genic, DNase hypersensitivy sites and open chromatin annotation were taken from the manifest files provided by Illumina and downloaded from the product support pages (MethylationEPIC_v-1-0_B2.csv).

### Genotyping and imputation

UKHLS samples were genotyped typed using the Illumina Infinium HumanCoreExome BeadChip Kit^®^ as previously described (12v1-0)(Prins et al. 2017). This array contains a set of >250,000 highly informative genome-wide tagging single nucleotide polymorphisms as well as a panel of functional (protein-altering) exonic markers, including a large proportion of low-frequency (MAF 1–5%) and rare (MAF <1%) variants. Genotype calling was performed with the gencall algorithm using GenomeStudio (Illumina Inc.). After selecting only the samples with matched DNAm data, variants were refiltered prior to imputation. PLINK (Purcell et al. 2007) was used to remove samples with >5% missing data. We also excluded SNPs characterized by >5% missing values, a Hardy-Weinberg equilibrium *P*-value < 0.001 and a minor allele frequency of <5%. To identify related samples, SNPs were LD pruned and the --genome command in PLINK was used to calculate the proportion of identity-by-descent for all pairs of samples; 58 pairs of related samples (PI_HAT > 0.2) were identified and one individual from each pair was randomly excluded to ensure the sample was independent. These data were then imputed using the 1000 genomes phase 3 version5 reference panel, SHAPEIT and minimac3 (Das et al. 2016). Best guess genotypes were called and variants filtered to those with minor allele frequency > 0.01 and INFO score > 0.8. As variants were named using their locations (“chr:pos”) and variant type (SNP/INDEL), duplicate variants were also excluded. Principal components were calculated from the imputed genotype data using GCTA(Yang et al. 2011). 16 samples appeared as outliers (defined as more than 2 standard deviations from the mean) in a scatterplot of the first two principal components and were excluded from subsequent genetic analyses. Principal components were then recalculated for inclusion as covariates in quantitative trait loci analyses. The imputed genetic variants were then filtered to exclude variants characterized by >5% missing values, a Hardy-Weinberg equilibrium *P*-value < 0.001, a minor allele frequency of <5% and a minimum of 5 observations in each genotype group. These genotype data are available on application through the European Genome-phenome Archive under accession EGAS00001001232 (https://www.ebi.ac.uk/ega/home).

### DNAm quantitative trait loci (mQTL)

Cross-hybridizing probes, probes with a common SNP (European population minor allele frequency > 0.01) within 10bp of the CpG site or single base extension (McCartney et al. 2016; Pidsley et al.

2016) and probes on the sex chromosomes were excluded from the QTL analysis. In addition, 977 substandard probes identified by Illumina were also excluded. We performed a genome-wide mQTL analysis; in total, 766,714 DNAm sites were tested against 5,210,475 genetic variants using the R package MatrixEQTL (Shabalin 2012). This package enables fast computation of QTLs by only saving those more significant than a pre-defined threshold (set to P = 1x10^-8^ for this analysis). An additive linear model was fitted to test if the number of alleles (coded 0,1,2) predicted DNAm at each site, including covariates for age, sex, six estimated cellular composition variables (B cells, CD8 T cells, CD4 T cells, monocytes, granulocytes, natural killer T cells)(Houseman et al. 2012; Koestler et al. 2013), two binary batch variables and the first ten principal components from the genotype data to control for ethnicity differences. A Bonferroni-corrected multiple testing threshold was used, set to genome-wide significance for GWAS divided by the number of DNAm sites tested (i.e. 5x10^-8^/766714 = 6.52x10^-14^). The *clump* command in PLINK(Purcell et al. 2007) was used to identify the number of independent associations for each DNAm site with more than 1 significant mQTL using the following parameters: --clump-p1 1e-8 --clump-p2 1e-8 --clump-r2 0.1 --clump-kb 250.

### Bayesian Co-localization

Taking all DNAm sites with at least 1 significant mQTL (P < 1x10^-10^), all pairs of DNAm sites located on the same chromosome and within 250kb of each other were tested for co-localization. As data for all SNPs (regardless of significance) are required for this analysis, first, the mQTL analysis was rerun for these DNAm sites to record all association statistics (p-value, regression coefficient and t-statistic to infer the standard error) for all SNPs within 500 kb of the DNAm site. Co-localization analysis was performed as previously described (Giambartolomei et al. 2014) using the R coloc package (http://cran.r-project.org/web/packages/coloc). From our mQTL results we inputted the regression coefficients, their variances and SNP minor allele frequencies, and the prior probabilities were left as their default values. This methodology quantifies the support across the results of each GWAS for 5 hypotheses by calculating the posterior probabilities, denoted as *PPi* for hypothesis *Hi*.

*H_0_: there exist no causal variants for either CpG site;*

*H_1_: there exists a causal variant for CpG_1_ only;*

*H_2_: there exists a causal variant for CpG_2_ only;*

*H_3_: there exist two distinct causal variants, one for each CpG;*

*H_4_: there exists a single causal variant common to both CpGs*.

### Summarised Mendelian Randomisation (SMR) analysis 1: identifying putative pleiotropic relationships between DNAm and complex traits

SMR analysis between DNAm and complex traits was performed as previously described (Zhu et al. 2016; Hannon et al. 2017) using the software downloaded from http://cnsgenomics.com/software/smr/. Publically available genome-wide association study results were downloaded from a range of sources and converted in the appropriate format for the SMR analysis. SNPs were renamed in 1000 genome format (chr:bp) to align with the mQTL output using dbSNP version 141 (where SNP locations for hg19 were not provided in the results file). Where allele frequency was not provided, it was taken from the European subset of the 1000 genomes (phase 3, version 5). Details for how each set of results was processed can be found in the **Supplementary Table 4**. Significant mQTL (P < 1x10^-10^) calculated in the UKHLS sample were used to identify genetic instruments for 126,457 DNAm sites included in the SMR analysis. The SMR test comprises of two steps, firstly a two sample Mendelian randomisation using the two-step least squares (2SLS) approach is applied using the effect size of the top cis-QTL SNP and its corresponding effect in the GWAS. The significance threshold for this part of test was set at 3.95x10^-7^, calculated using the Bonferroni correction method and adjusting for the number of DNAm sites tested (0.05/126,457). The second step tests for heterogeneity of effects using alternative SNPs as the instrumental variable, based on the theory that if both DNAm and the GWAS trait are associated with the same causal variant, the choice of SNP is irrelevant, whereas if they were associated with different causal variants, the differing linkage disequilibrium relationships between the instruments and each causal variant would lead to variation in the estimated effect between the trait and DNAm. Non-significant heterogeneity (HEIDI P > 0.05) indicates that there is a pleiotropic effect on a GWAS trait and DNAm. This approach was also applied using publically available eQTL data from (Westra et al. 2013), in this analysis significant pleotropic associations between gene expression and complex traits were selected as those with SMR P < 8.38x10^-6^ (corrected for 5,966 gene expression probes tested) and HEIDI *P* > 0.05.

### Summarised Mendelian Randomisation (SMR) analysis 2: identifying putative pleiotropic relationships between DNAm and gene expression

A second application of the SMR analysis was used to identify pleiotropic relationships between DNAm and gene expression. Gene expression quantitative trait loci (eQTL) results from the Westera eQTL study (Westra et al. 2013) were downloaded from http://cnsgenomics.com/software/smr/download.html. SNP ids were converted to 1000 genomes format (in order to match the mQTL output) and SNP frequencies were taken from the European subset 1000 genomes (phase 3, version 5). These data included eQTL at 5,966 probes. All pairs of CpG and genes where tested providing that a) the CpG had a significant mQTL (P < 1x10^-10^), b) the gene had a significant eQTL (P < 5x10^-8^) and c) there was a common genetic variant tested within 500 kilobases of the gene expression probe and DNAm site. In total 488,342 pairs of DNAm sites and gene expression transcripts were tested hence the significance threshold for the first stage of the SMR test was set to P < 1.02x10^-7^ after applying a Bonferroni correction for the number of tests. Consistent with all other SMR analyses in this manuscript, a non-significant heterogeneity test (HEIDI) > 0.05) in step 2 of the SMR analysis was used to classify pleiotropic relationships from artefacts of linkage disequilibrium.

### Enrichment analyses

DNAm sites were annotated to genes and CpG islands using the manifest file provided by Illumina. Gene annotation is based on the UCSC RefGene database. Chi-square tests were used to test for different distribution of genic annotation categories using all tested DNAm sites as the background.

## DATA ACCESS

Individual level DNA methylation and genetic data are available on application through the European Genome-phenome Archive under accession EGAS00001001232 (https://www.ebi.ac.uk/ega/home). Specific details can be found here (https://www.understandingsociety.ac.uk/about/health/data). Phenotype linked to DNA methylation data are available through application to the METADAC (www.metadac.ac.uk). Summary statistics for all Bonferroni significant DNA methylation quantitative trait loci are available for download from http://www.epigenomicslab.com/online-data-resources/. Many of the results included in this manuscript are can be explored through our interactive web application also available at http://www.epigenomicslab.com/online-data-resources/. Analysis scripts used in this manuscript are available on https://github.com/ejh243/UKHLS_mQTL.git.

## ACKNOWLEDGMENTS

We acknowledge the Wellcome Trust Sanger Institute and Ele Zeggini for generating the genotype data. Both genotyping and DNA methylation in UKHLS were funded through enhancements to the Economic and Social Research council (ESRC) grants ES/K005146/1 and ES/N00812X/1. AH, MS, YB are supported by the ESRC (ES/M008592/1). MK is supported by the University of Essex and ESRC (RES-596-28-0001). EH, JM and LS time on this project was supported by MRC grant K013807. Analysis was facilitated by access to the Genome high performance computing cluster at the University Of Essex School Of Biological Sciences.

## SUPPLEMENTAL DATA

### SupplementaryFile1.pdf

**Manhattan plots of Summary data-based Mendelian Randomisation (SMR) tests for pleiotropic effects between 63 complex traits and DNA methylation.** Shown on the y-axis of each plot is the – log10 P-value from the SMR analysis using DNA methylation quantitative trait loci (mQTL) generated from whole blood. Each point represents an SMR test for a particular DNA methylation site. The red horizontal line represents the genome-wide multiple testing significance threshold (P < 6.42 x 10^-7^); green points highlight the significant SMR tests which are not characterized by significant heterogeneity (i.e. P > 0.05), indicating pleiotropic relationships between that trait and either DNA methylation.

**Supplementary Table 1.** Summary of DNA methylation quantitative trait loci (mQTL) analysis in the UKHLS cohort (n = 1,111) for A) all mQTL, B) mQTL classified as *cis* (where the distance between the genetic variant and DNA methylation site < 500kb), and C) mQTL classified as *trans*.

**Supplementary Table 2.** Summary of the additional mQTL associations identified using the Illumina EPIC array not detected using the Illumina 450K array.

**Supplementary Table 3.** Frequency of DNAm sites in A) genic feature and B) CpG Island feature annotation categories.

**Supplementary Table 4.** All DNAm site pairs characterized by a higher posterior probability for both DNAm sites being associated with the same causal variant as determined by Bayesian co-localisation analysis (PP3 + PP4 > 0.99 & PP4/PP3 > 1).

**Supplementary Table 5.** Details of all publically available GWAS results included in Summary data-based Mendelian Randomization (SMR) analysis.

**Supplementary Table 6.** Results from SMR analysis highlighting all significant associations (P < 3.95x10^-7^) between a complex trait and DNA methylation.

**Supplementary Table 7.** Overlap of SMR analysis results using DNA methylation quantitative trait loci (mQTL) and gene expression quantitative trait loci (eQTL) where a complex trait is pleiotropically associated with both DNA methylation and gene expression at the same gene.

**Supplementary Table 8.** Significant pleiotropic associations between gene expression and DNA methylation (where SMR P < 1.02x10-7 & HEIDI P > 0.05).

**Supplementary Table 9.** Frequency of DNAm sites associated with gene expression in gene feature annotation categories.

**Supplementary Figure 1:** Overview of the study providing a schematic of our analytical plan and the datasets used in analyses.

**Supplementary Figure 2:** Genic distribution of novel associations between genetic variation and DNA methylation using the Illumina EPIC array.

**Supplementary Figure 3:** Additional content on the Illumina EPIC array enable the identification of novel DNA methylation quantitative trait loci (mQTL) associations.

**Supplementary Figure 4:** Frequency distribution of DNA methylation quantitative trait loci (mQTL) SNPs and their associated DNA methylation sites.

**Supplementary Figure 5:** Frequency distribution of DNA methylation quantitative trait loci (mQTL) SNPs and associated DNA methylation sites after filtering for independent associations.

**Supplementary Figure 6:** The distribution of effect sizes across all Bonferroni significant DNA methylation quantitative trait loci (mQTL).

**Supplementary Figure 7:** The significance and effect size of *cis* mQTL associations increases as the distance between the genetic variant and DNAm site decreases.

**Supplementary Figure 8:** Genic distribution of DNA methylation sites associated with mQTL variation.

**Supplementary Figure 9:** Distribution of DNA methylation sites associated with mQTL variation in CpG island features.

**Supplementary Figure 10:** DNA methylation sites associated with common mQTL variants are more strongly influenced by additive genetic variation.

**Supplementary Figure 11:** Distribution of correlation coefficients between pairs of DNAm sites with shared genetic effects.

**Supplementary Figure 12:** Distribution of correlation coefficients between pairs of DNAm sites with shared genetic effects, split by genic feature annotation.

**Supplementary Figure 13:** Replication of significant pleiotropic associations across two independent datasets.

**Supplementary Figure 14:** Flowchart showing the number of gene expression probes, DNAm sites and pairs significant at each stage of the analysis.

**Supplementary Figure 15:** Genomic distribution of DNA methylation sites pleiotropically associated with gene expression.

**Supplementary Figure 16:** Distribution of estimated effect of pleiotropic associations between DNA methylation and gene expression, stratified by genic location of DNA methylation site.

